# Visualizing tumour self-homing with magnetic particle imaging

**DOI:** 10.1101/2020.02.17.953232

**Authors:** Katie M Parkins, Kierstin P Melo, John A Ronald, Paula J Foster

## Abstract

Due to their innate tumour homing capabilities, in recent years, CTCs have been engineered to express therapeutic genes for targeted treatment of primary and metastatic lesions. Additionally, previous studies have incorporated optical or PET imaging reporter genes to enable noninvasive monitoring of therapeutic CTCs in preclinical tumour models. Here, we demonstrate for the first time, the ability of magnetic particle imaging (MPI) to sensitively detect systemically administered iron-labeled CTCs and to visualize tumour self-homing in a murine model of human breast cancer.

## Introduction

Tumour self-homing describes a phenomenon where circulating tumour cells (CTCs) that have shed from a primary tumour into the circulation, can return to grow at their original tumour site. This concept was first described by Norton and Massagué in 2006 and is thought to be driven by both a leaky vasculature that permits CTC recruitment, as well as a permissive tumour microenvironment that promotes CTC survival and growth^1,2^. Due to their tumour targeting capabilities, in recent years, self-homing CTCs have been repurposed as delivery vehicles for anti-cancer therapeutics. This has included the delivery of oncolytic viruses, pro-drug activatable suicide genes, and transgenes that alter the tumour microenvironment^3-9^. This strategy has shown exciting progress towards treating primary tumours, single organ metastases and most recently, multi-organ metastases however, further refinement is needed in order to optimize self-homing CTCs for potential clinical translation. Tools that enable the fate of systemically administered CTCs to be noninvasively monitored over time would provide valuable information about the kinetics and efficiency of CTC infiltration into tumours, their proliferation and persistence over time, any unwanted off-tumour accumulation, as well as information to better understand therapeutic response in individual subjects.

Cellular imaging can be used to noninvasively study a specific cell population or cellular process *in vivo.* Previous studies have used imaging reporter genes, namely bioluminescence imaging (BLI)^4,9^ and positron emission tomography (PET) reporters^7,8^, to track CTCs in preclinical models. For instance, in 2007, Power and colleagues used dual-enzyme BLI, a commonly used optical modality, to noninvasively monitor both the carrier cells (i.e., cancer cells) and their viral payload *in vivo*. By using this system, they demonstrate independent monitoring of cell vehicle distribution and virus associated luciferase enzymes. Our group has recently used dual BLI to visualize spontaneous whole-body breast cancer metastases expressing *Renilla* luciferase, and systemically administered theranostic CTCs expressing *Firefly* luciferase. Additionally, as a step towards tracking CTCs in patients, Reinshagen et al., recently demonstrated that the PET reporter gene, herpes simplex virus-thymidine kinase (HSV-TK) can be used to monitor the fate of therapeutic CTCs derived from glioblastoma.

An alternative method for tracking cells is to pre-label them with imaging probes prior to transplantation into the body^10-16^. These techniques are complementary to reporter gene techniques and often used for short term tracking of the initial arrest of cells at a target site, due to probe dilution during cell division. An advantage of probe-based tracking is that it does not require genetic engineering of cells, which may be more feasible for translational purposes. Moreover, probe-based cell tracking is often much more sensitive to low numbers of cells compared to reporter gene technologies due to the fact that large amounts of probe can be concentrated into each cell. Our group has previously shown MRI of cells loaded with superparamagnetic iron oxide nanoparticles (SPIONs) with single cell sensitivity in various mouse models^13,14,16^. We and others have used this extensively to track cell types such as cancer cells, stem cells and immune cells. However, some limitations of SPION-based MRI are that SPIONs create a loss of signal and quantitation of signal loss and the number of iron-labelled cells in a particular region is challenging^11^. Magnetic particle imaging (MPI) is an emerging imaging technique that sensitively and specifically detects superparamagnetic iron oxide nanoparticles (SPIONs)^17-19^. In MPI, SPIONs result in positive signal and the signal strength is linearly proportional to the number of SPIONs, which allows for truly quantitative imaging. A few groups have started to explore the potential of MPI as a novel cell tracking technology with various SPIONs (e.g., ferumoxytol or ferucarbotran)^18-21^. However, very few groups have explored the use of MPI to track the biodistribution of systemically administered SPION-labeled cells. In this work, we monitor the fate of SPION-labeled experimental CTCs in tumour bearing mice and demonstrate for the first time, the visualization of tumour self-homing in a mouse model of breast cancer with high sensitivity MPI.

## Methods

### Cell labeling

MDA-MB-231 and MDA-MB-231BR-eGFP cells were each maintained in Dulbecco’s modified eagle media (DMEM) containing 10% fetal bovine serum (FBS) at 37 °C and 5 % CO_2._ For cell labeling, 2 × 10^6^ adherent cells were incubated with 25 μg Fe/mL micron-sized paramagnetic iron-oxide (MPIO) beads (0.9 μm in diameter, 63% magnetite, labeled with Flash Red; Bangs Laboratory, Fishers, IN, USA) for 24 hours. Cells were washed three times with Hanks balanced salt solution (HBSS) and then trypsinized with 0.25% Trypsin-EDTA. The cells were then collected and thoroughly washed three more times with HBSS to remove unincorporated MPIO before cell injection and *in vitro* evaluation. Cell labeling had no effect on cell viability, and labeling efficiency was assessed by Perl’s Prussian blue (PPB) staining.

### Animal Model

The animals were cared for in accordance with the standards of the Canadian Council on Animal Care, and under an approved protocol (2015-0558) of the University of Western Ontario’s Council on Animal Care. A primary mammary fat pad (MFP) tumour was generated by injecting 1 × 10^6^ MDA-MB-231 cells into the lower right MFP of 4 female NOD scid gamma (NSG) mice (6-7 weeks old; Charles River Laboratories, Wilmington, MA, USA). Cells were suspended in 0.05 mL of HBSS per injection. After 41 days of primary tumour growth, mice received an intracardiac injection of 5 × 10^5^ MPIO labeled MDA-MB-231BR-eGFP cells into the left ventricle. Cells were suspended in 0.1 mL of HBSS and image guided injections were performed using a Vevo 2100 ultrasound system (VisualSonics Inc., Toronto, ON, CAN). Tumour volume was manually measured with calipers in two perpendicular dimensions, and the volume was estimated using the following formula = 0.52 (width)^2^(length), to approximate the volume of an ellipsoid (mm^3^)^22,23^.

### MPI Acquisition

Full body MPI images of tumour-bearing mice were acquired 72 hours following intracardiac injection of MPIO-labeled CTCs (day 44). Images were collected on a Momentum™ scanner (Magnetic Insight Inc., Alameda, CA, USA) using the 3D high sensitivity scan mode. In this mode, tomographic images were acquired using a 3 T/m gradient, 35 projections and a FOV 12 × 6 × 6 cm, for a total scan time ∼1 hour per mouse. Mice were anesthetized with 2% isoflurane in 100% oxygen during these scans. 3D high sensitivity images of *ex vivo* tumours were acquired using the same parameters.

### MPI Calibration and Signal Quantification

To generate a calibration curve, a phantom was made with 1 μL aliquots of MPIO beads and imaged using the same parameters as *in vivo* images. The following samples of iron content were tested: 0.07 μg, 0.105 μg, 0.14 μg, 0.21 μg, 0.28 μg, 0.7 μg, 1.05 μg, 1.4 μg, 2.1 μg, and 2.8 μg of iron. Images were analyzed utilizing Horos imaging software (Annapolis, MD USA). To calculate the total MPI signal in each image set, signal intensities were set to full dynamic range to best represent the full range of signal in a specific region of interest (ROI; i.e. calibration samples or tumours), prior to manually outlining the signal. The signal intensities from the *in vivo* images had to be adjusted to visualize tumour signal, decreasing window level and width to compensate for the high liver signal (max = 0.133, min = 0.01). Areas of interest from *in vivo* 3D images were manually outlined, slice by slice, creating a 3D volume. These ROIs (determined from tumour signal) were copied and pasted onto the contralateral MFP of the same mouse to assure an equivalent volume of control tissue was used for quantification of signal. Total MPI signal was calculated by <u>*mean signal x volume(mm*^*3*^)</u>. Total MPI signal was plotted against iron content to derive calibration lines and quantify iron content in tumours and contralateral MFPs.

### MRI Acquisition

MRI scans were performed on a 3 T MR750 clinical scanner (General Electric) equipped with a custom-built, insertable gradient coil and mouse brain solenoidal radiofrequency coil^13,16^. Tumour samples were tightly placed in MR compatible tubes to avoid motion artifacts. Images were acquired using a balanced Steady State Free Precession (bSSFP) imaging sequence [Fast Imaging Employing Steady State Acquisition (FIESTA) on the GE system] which has been previously optimized for iron detection^24^. The scan parameters were: repetition time (TR) = 7ms, echo time (TE) = 3.5 ms, bandwidth (BW) = 31.25kHz, flip angle (FA) = 35 degrees, averages (NEX) = 2, phase cycles = 4, matrix = 200 × 200. Total scan time was approximately 25 minutes per sample.

### Histology and immunohistochemistry

Following imaging, all mice were sacrificed by isoflurane overdose and perfused with 4% paraformaldehyde. Tumours were excised and placed in paraformaldehyde for an additional 24 hours. Fixed tissue was processed, paraffin embedded and cut into 10 *μ*m sections. Select sections were stained for iron with Perl’s Prussian blue (PPB) or stained for GFP by immunohistochemistry.

### Statistical Analysis

Statistics were calculated using GraphPad Prism 7 Software. Pearson’s rank correlation was used to determine the relationship between total MPI signal and iron content. *In vivo* data was expressed as mean ± SEM and analyzed by a Student’s t test. Differences were considered statistically significant at *p < 0.05.

## Results

### In vitro studies

Figure 1A shows the MDA-MB-232BR-GFP cell line was efficiently labeled with MPIO. The Perl’s Prussian blue stain shows intracellular iron in blue within the breast cancer cells that appear pink. We acquired images of samples with known iron content to determine the relationship between MPI signal and iron content (Figure 1B). Samples were separated by 2cm on the MPI bed to allow for imaging of 5 samples per scan. A strong relationship between iron content and MPI signal was observed (R^2^= 0.974, p < 0.0001) (Figure 1C). These calibration curves were used to quantify iron content of MPIO labeled MDA-MB-231BR cells in *in vivo* and *ex vivo* images.

**Figure 1:**
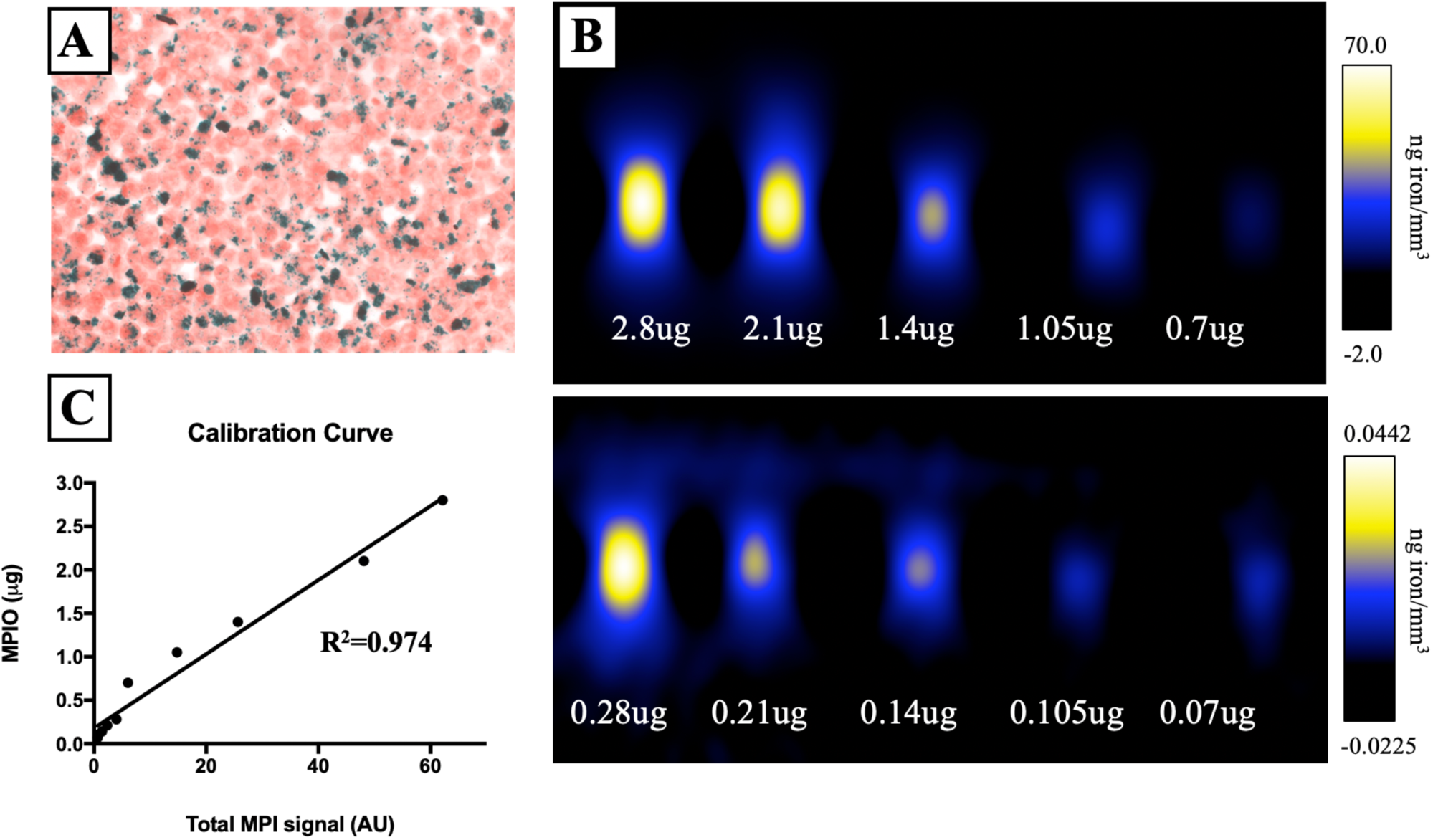
Experimental CTCs were efficiently labeled with iron as seen with PPB staining (iron=blue) (A). Images of samples with known iron content were used to generate a calibration curve for quantification purposes (B). A strong relationship between iron content and MPI signal was observed (R^2^= 0.974, p < 0.0001) (C).

### In vivo studies

Figure 2A shows MPI images of tumour bearing mice 72 hours after receiving 5 × 10^5^ MPIO labeled MDA-MB-231BR cells. These images were scaled to display the full dynamic range of signal across all mice. MPI signal can be clearly visualized in the lower right MFP tumour. Signal was also detected in lungs of some of the mice, likely due to iron labeled cells that were trapped in the lungs following intracardiac injection, as well as, signal in the abdomen, presumably due to iron content in the mouse feed. The distribution of MPI signal throughout the body can be visualized by scrolling through the complete 3D dataset (Video file 1). We found that iron content in the lower right MFP (tumour bearing) (M= 0.798 ± 0.184μg) was significantly higher than in the contralateral MFP (M= 0.318 ± 0.044μg). The average tumour volume measured by calipers was 230.6 ± 42.48mm^3^.

**Figure 2:**
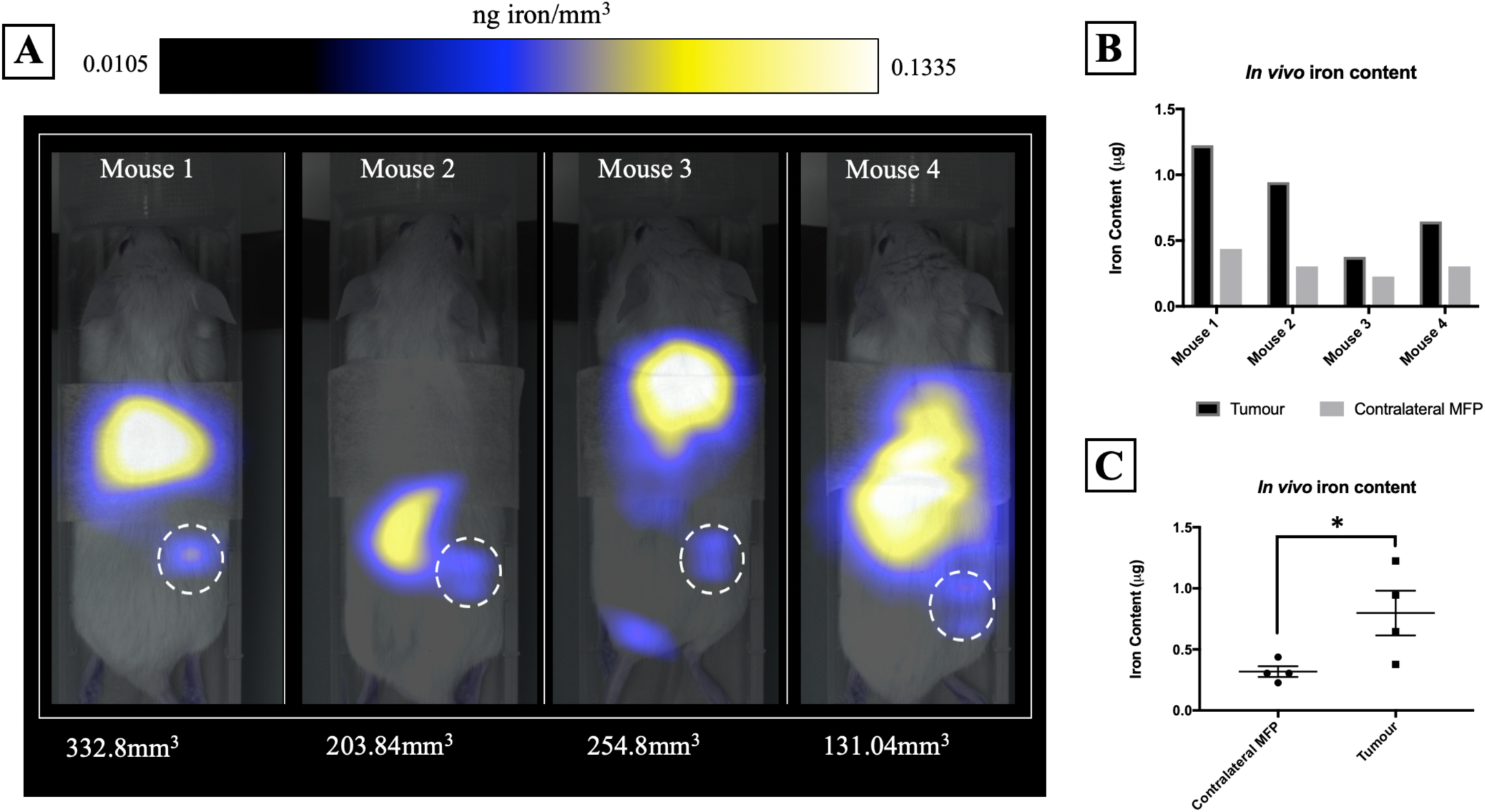
MPI signal, representing iron-labeled experimental CTCs, was detected in the lower right MFP tumour of all mice (A). MFP tumour burden (mm^3^) measured by calipers is shown directly below each mouse. Iron content in the lower right MFP (tumour bearing) (M= 0.798 ± 0.184μg) was significantly higher than in the contralateral MFP (M= 0.318 ± 0.044μg) (B/C).

### Ex vivo studies

MPIO labeled MDA-MB-231BR cells were also visualized within MFP tumours using *ex vivo* imaging (Figure 3). In MPI images, the iron distribution of 3 different tumour samples of varying sizes were visualized (Figure 3A/B). The average iron content per tumour was 1.48 ± 0.24μg (Figure 3C). Iron labeled cells were also visualized as regions of signal void throughout tumours using iron-sensitive *ex vivo* MRI (Figure 3D).

**Figure 3:**
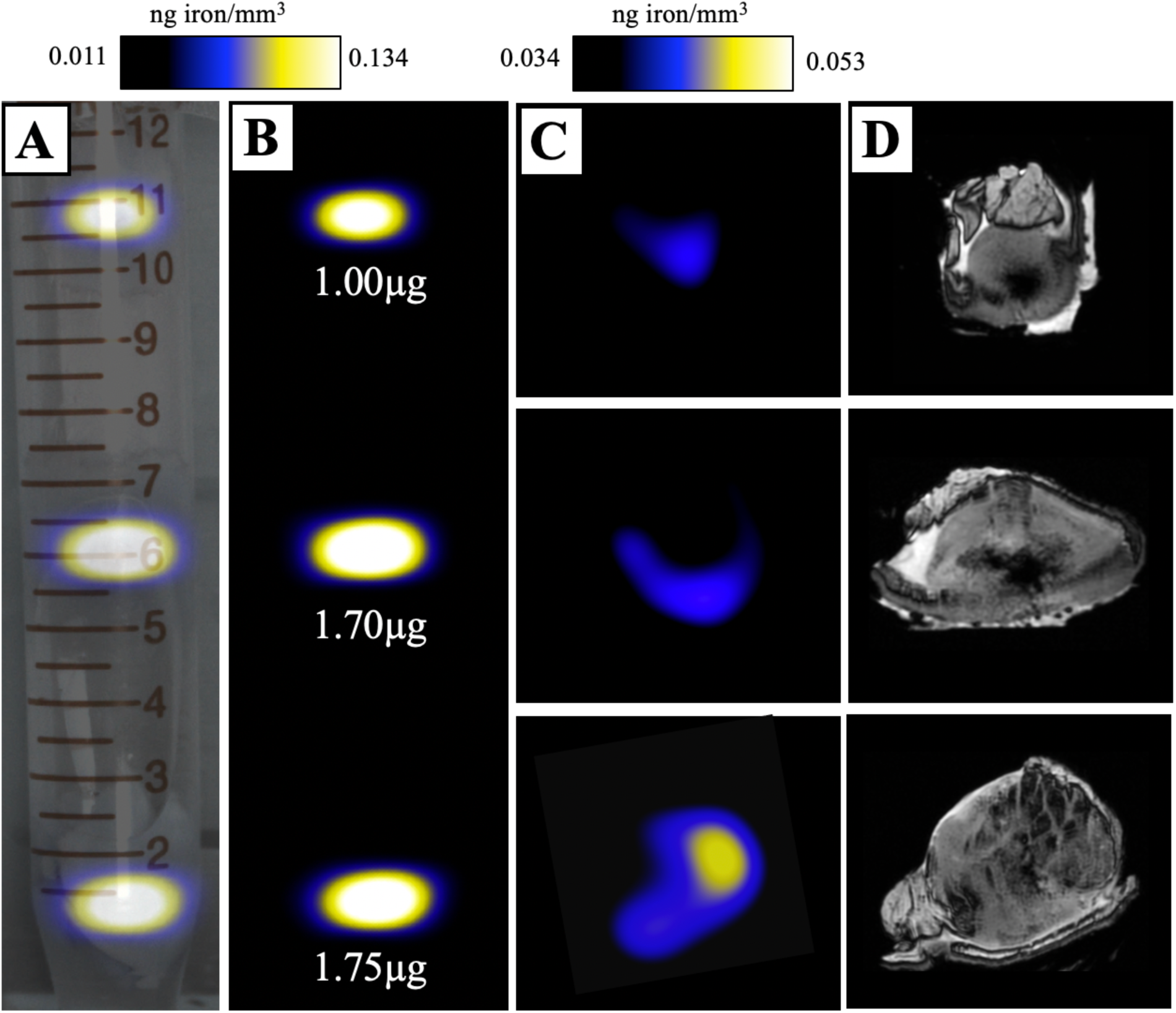
Ex vivo tumour samples were placed in a 15mL falcon tube and imaged with MPI and MRI. A maximum intensity projection of MPI signal was overlaid onto a brightfield image for anatomical context (A/B). Individual slices from the 3D MPI data set were qualitatively assessed for correspondence with areas of signal void in MRI (C/D).

### Microscopy and Immunohistochemistry

Mice were sacrificed 44 days after MFP cell injection. Iron labeled cancer cells were visualized within the MFP tumour using PPB staining (Figure 4A). Immunostaining of adjacent sections demonstrated that the location of GFP positive cells corresponded with the location of iron (Figure 4B). This suggests the MPI and MRI signal we are seeing in iron-based imaging techniques is from iron-labeled, GFP expressing CTCs throughout the tumour.

**Figure 4:**
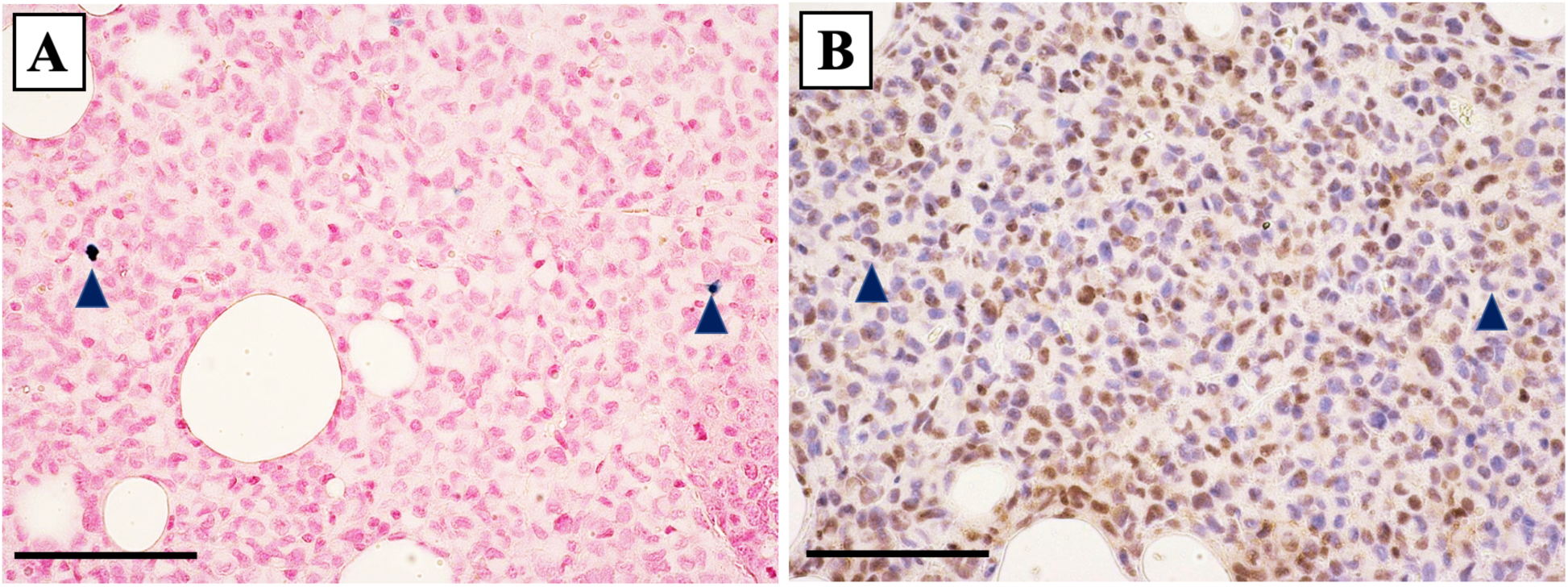
Iron labeled CTCs (blue) were visualized within the MFP tumour using PPB staining (A). Immunostaining of adjacent tumour sections confirmed the presence of GFP positive cells (brown) including some that correspond with the location of iron (B). (40X magnification with scale bar 100µm)

## Discussion

A number of different cell-based vectors have previously been developed for the targeted delivery of anti-cancer therapeutics to primary and metastatic lesions^25-31^. While some cell types have shown promise due to their innate homing capabilities (i.e. Immune cells and stem cells), new strategies to overcome intratumoural immunological barriers and on-target off-tumour effects are urgently needed. CTCs may represent a novel cell-based platform for therapy whereby, they are highly efficient at homing to established tumour sites, can be readily engineered and expanded *ex vivo*, and could be generated from an individual patient’s tumour to avoid an unwanted immune response. Our group and others have shown exciting progress towards the development of self-homing CTCs for the treatment of primary and metastatic lesions however, further study is warranted to optimize CTCs for clinical translation^3-9^. Here, we demonstrate for the first time, the ability of MPI to sensitively track SPION-loaded experimental CTCs and to visualize tumour self-homing in a murine model of human breast cancer.

In this work, we visualized iron-labeled MDA-MB-231BR-eGFP cells that had migrated to an MDA-MB-231 primary tumour, which we hypothesized may be due to a well-established tumour microenvironment. Previous work by Kim *et al*., has shown that an MDA-MB-231 tumour can produce chemoattractants, IL-6 and IL-8, that are capable of actively recruiting shed CTCs^32^. Vilalta and colleagues have also shown in the murine 4T1 breast cancer model that irradiation of tumours induces the expression of granulocyte-macrophage colony stimulating factor (GM-CSF) that can act to enhance the recruitment of CTCs^33^. Their evidence suggests radiation induced self-homing may act as a potential mechanism for cancer recurrence in the clinic. Future work will look to investigate how the production of these cytokines relates to the amount of CTC self-homing, and whether we can visualize these differences over time with MPI.

In our model, the primary MDA-MB-231 MFP tumour had 41 days to grow prior to the administration of experimental CTCs. Despite observing a range in primary tumour size across mice, a clear relationship between MPI signal, representing the amount of CTC homing, and the size of the MFP tumour was not observed. Furthermore, the mouse having the smallest measured tumour burden (approximately 133mm^3^) had MPI signal far above our estimated detection threshold, suggesting MPI may have the capability to detect much fewer CTCs than shown in this study. Future investigation should look to decrease tumour burden as well as alter the number of CTCs that are administered to better determine the sensitivity of MPI for detecting and longitudinally monitoring therapeutic CTCs *in vivo*. Additionally, the ratio of tumour cells to therapeutic CTCs needed for a therapeutic effect will be important when moving towards the treatment of whole-body metastases.

Our group and others have previously applied cellular MRI to track various iron-loaded cell types *in vivo* including immune cells, stem cells, cancer cells and pancreatic islets. Previous work has shown that labeling cancer cells with iron does not cause significant changes in cell viability, proliferation, apoptosis or metastatic efficiency making it an ideal probe for clinically relevant imaging studies^16,34^. However, iron-based cell tracking techniques provide a limited imaging window as the probe gets diluted through cancer cell division^15^. In this work, we performed MPI at 72 hours following systemic injection of iron-labeled experimental CTCs. While we saw substantial MPI signal in all of the tumours, signal could be further enhanced by optimizing the timeline to visualize the maximum amount of self-homing prior to probe dilution. Furthermore, we observed MPI signal in the abdomen of mice as a result of having iron in the mouse feed. This was not detrimental to our study since the tumour bearing MFP is far enough away from the stomach to avoid signal overlap however, an iron-free diet should be considered for future studies that may include disease within that region.

## Conclusions

In this work, we employed high sensitivity MPI to noninvasively visualize SPION-loaded experimental CTCs in tumour bearing mice. Further, we demonstrate a clinically relevant model of tumour self-homing whereby, SPION-labeled CTCs migrated, and were detected at the established MFP tumour site 72 hours following systemic administration. Future work will look to further build MPI as a valuable tool for visualizing whole-body tumour self-homing, which will be extremely valuable in understanding the fate of therapeutic CTCs and the potential mechanisms driving tumour self-homing throughout the body.

## Acknowledgements

The authors would like to acknowledge funding from the Canadian Foundation for Innovation.

## Conflicts of Interest

There are no conflicts of interest to declare.

## References

1. Norton L, Massague J. Is cancer a disease of self-seeding? Nat Med 2006;12:875–878.

2. Carmeliet P, Jain RK. Angiogenesis in cancer and other diseases. Nature. 2000;407:249–257.

3. Z. Raykov, G. Balboni, M. Aprahamian, J. Rommelaere, Carrier cell-mediated delivery of oncolytic parvoviruses for targeting metastases. Int. J. Cancer 109, 742–749 (2004).

4. A. T. Power, J. Wang, T. J. Falls, J. M. Paterson, K. A. Parato, B. D. Lichty, D. F. Stojdl, P. A. Forsyth, H. Atkins, J. C. Bell, Carrier cell-based delivery of an oncolytic virus circumvents antiviral immunity. Mol. Ther. 15, 123–130 (2007).

5. S. M. Freeman, C. N. Abboud, K. A. Whartenby, C. H. Packman, D. S. Koeplin, F. L. Moolten, G. N. Abraham, The “bystander effect”: Tumor regression when a fraction of the tumor mass is genetically modified. Cancer Res. 53, 5274–5283 (1993).

6. J. Garcia-Castro, J. Mart. nez-Palacio, R. Lillo, F. Garc.a-S. nchez, R. Alemany, L. Madero, J. A. Bueren, M. Ram. rez, Tumor cells as cellular vehicles to deliver gene therapies to metastatic tumors. Cancer Gene Ther. 12, 341–349 (2005).

7. E. Dondossola, A. S. Dobroff, S. Marchi., M. Card.-Vila, H. Hosoya, S. K. Libutti, A. Corti,R. L. Sidman, W. Arap, R. Pasqualini, Self-targeting of TNF-releasing cancer cells in preclinical models of primary and metastatic tumors. Proc. Natl. Acad. Sci. U.S.A. 113, 2223–2228 (2016).

8. Reinshagen, C., Bhere, D., Choi, S.H., Hutten, S., Nesterenko, I., Wakimoto, H., Le Roux, E., Rizvi, A., Du, W., Minicucci, C. and Shah, K., 2018. CRISPR-enhanced engineering of therapy-sensitive cancer cells for self-targeting of primary and metastatic tumors. Science translational medicine, 10(449), p.eaao3240.

9. Parkins KM, Dubois VP, Kelly JJ, Chen Y, Foster PJ, Ronald JA. “Engineering self-homing circulating tumor cells as novel cancer theranostics”. Pre-print: bioRxiv 746685; doi: https://doi.org/10.1101/746685

10. Gaudet JM, Hamilton AM, Chen Y, Fox MS, Foster PJ. Application of dual 19F and iron cellular MRI agents to track the infiltration of immune cells to the site of a rejected stem cell transplant. Magn Reson Med. 2017;78(2):713–20.

11. Makela A V., Murrell DH, Parkins KM, Kara J, Gaudet JM, Foster PJ. Cellular imaging with MRI. Top Magn Reson Imaging. 2016;25(5):177–86.

12. Khurana A, Nejadnik H, Gawande R, Lin G, Lee S, Messing S, Castaneda R, Derugin N, Pisani L, Lue TF, Daldrup-Link HE. Intravenous ferumoxytol allows noninvasive MR imaging monitoring of macrophage migration into stem cell transplants. Radiology. 2012;264(3):803–11.

13. Heyn C, Bowen C V., Rutt BK, Foster PJ. Detection threshold of single SPIO-labeled cells with FIESTA. Magn Reson Med. 2005; 53(2):312

14. Parkins KM, Hamilton AM, Makela AV, Chen Y, Foster PJ, Ronald JA. “A multimodality imaging model to track viable cancer cells from single arrest to metastasis” Scientific Reports. October 2016.

15. Economopoulos, V., Chen, Y., McFadden, C., Foster, PJ. MRI Detection of Nonproliferative Tumor Cells in Lymph Node Metastases Using Iron Oxide Particles in a Mouse Model of Breast Cancer. Transl Oncol. 6:347–54. (2013).

16. Heyn, C., Ronald, JA., Ramadan, SS., Snir, JA., Barry, AM., MacKenzie, LT., et al. In vivo MRI of cancer cell fate at the single-cell level in a mouse model of breast cancer metastasis to the brain. Magn Reson Med. 56:1001–10. (2006).

17. Bulte JWM. Superparamagnetic iron oxides as MPI tracers: A primer and review of early applications. Adv Drug Deliv Rev [Internet]. 2019;138:293–301.

18. Nejadnik H, Pandit P, Lenkov O, Lahiji AP, Yerneni K, Daldrup-Link HE. Ferumoxytol Can Be Used for Quantitative Magnetic Particle Imaging of Transplanted Stem Cells. Molecular Imaging and Biology. 2018; 21(3):465–472.

19. Bulte JWM, Walczak P, Janowski M, Krishnan KM, Arami H, Halkola A, Gleich B, Rahmer J. Quantitative “Hot-Spot” Imaging of Transplanted Stem Cells Using Superparamagnetic Tracers and Magnetic Particle Imaging. Tomography. 2015;1(2):91–7

20. Sehl, O. C., Makela, A. V., Hamilton, A. M., & Foster, P. J. (2019). Trimodal Cell Tracking In Vivo: Combining Iron-and Fluorine-Based Magnetic Resonance Imaging with Magnetic Particle Imaging to Monitor the Delivery of Mesenchymal Stem Cells and the Ensuing Inflammation. Tomography, 5(4), 367.

21. Gaudet, J. M., Makela, A. V., & Foster, P. J. (2019). Non-invasive detection and quantification of tumor-associated macrophage density with magnetic particle imaging.

22. Euhus, D. M., Hudd, C., LaRegina, M. C. & Johnson, F. E. Tumor measurement in the nude mouse. J Surg Oncol. 31, 229–234 (1986). 21.

23. Tomayko, M. M. & Reynolds, C. P. Determination of subcutaneous tumor size in athymic (nude) mice. Cancer Chemother. Pharmacol. 24, 148–154 (1989).

24. Ribot, E. et al. *In vivo* single scan detection of both iron-labeled cells and breast cancer metastases in the mouse brain using balanced steady-state free precession imaging at 1.5 T. J. Magn Reson Imaging. 34, 231–238 (2011).

25. M. Mimeault, R. Hauke, S. K. Batra, Stem cells: A revolution in therapeutics—Recent advances in stem cell biology and their therapeutic applications in regenerative medicine and cancer therapies. Clin. Pharmacol. Ther. 82, 252–264 (2007).

26. J. Ankrum, J.M. Karp, Mesenchymal stem cell therapy: two steps forward, one step back, Trends Mol. Med. 16 (2010) 203–209.

27. K. Aboody, et al., Neural stem cells display extensive tropism for pathology in adult brain: evidence from intracranial gliomas, Proc. Natl. Acad. Sci. U. S. A. 97 (2001) 12846–12851.

28. K.I. Park, et al., Neural stem cells may be uniquely suited for combined gene therapy and cell replacement: evidence from engraftment of Neurotrophin-3-expressing stem cells in hypoxic–ischemic brain injury, Exp. Neurol. 199 (2006) 179–190.

29. M. Alieva, et al., Glioblastoma therapy with cytotoxic mesenchymal stromal cells optimized by bioluminescence imaging of tumor and therapeutic cell response, PLoS One 7 (2012) 1–11.

30. J.R. Bagó, K.T. Sheets, S.D. Hingtgen, Neural stem cell therapy for cancer, Methods. 99 (2016) 37–43.

31. K. Song, N. Benhaga, R.L. Anderson, R. Khosravi-Far, Transduction of tumor necrosis factor-related apoptosis-inducing ligand into hematopoietic cells leads to inhibition of syngeneic tumor growth in vivo, Cancer Res. 66 (2006) 6304–6311.

32. Kim, M. Y., Oskarsson, T., Acharyya, S., Nguyen, D. X., Zhang, X. H. F., Norton, L., & Massagué, J. (2009). Tumor self-seeding by circulating cancer cells. Cell, 139(7), 1315–1326.

33. Vilalta, M., Rafat, M., Giaccia, A. J., & Graves, E. E. (2014). Recruitment of circulating breast cancer cells is stimulated by radiotherapy. Cell reports, 8(2), 402–409.

34. Rohani, R., de Chickera, SN., Willert, C., Chen, Y., Dekaban, GA., Foster, PJ. In vivo cellular MRI of dendritic cell migration using micrometer-sized iron oxide (MPIO) particles. Mol Imaging Biol. 13:679–94. (2011).

